# Decoding trans-saccadic prediction error

**DOI:** 10.1101/2022.03.21.485172

**Authors:** Louise Catheryne Barne, Jonathan Giordano, Thérèse Collins, Andrea Desantis

**Author notes:** **Corresponding author email address** Louise Catheryne Barne. Co-senior authors (Thérèse Collins, Andrea Desantis).

## Abstract

We are constantly sampling our environment by moving our eyes, but our subjective experience of the world is stable and constant. Stimulus displacement during or shortly after a saccade often goes unnoticed, a phenomenon called the saccadic suppression of displacement. Although we fail to notice such displacements, our oculomotor system computes the prediction errors and adequately adjusts the gaze and future saccadic execution, a phenomenon known as saccadic adaptation. In the present study, we aimed to find a brain signature of the trans-saccadic prediction error that informs the motor system but not explicit perception. We asked participants (either sex) to report whether a visual target was displaced during a saccade while recording electroencephalography (EEG). Using multivariate pattern analysis, we were able to differentiate displacements from no displacements, even when participants failed to report the displacement. In other words, we found that trans-saccadic prediction error is represented in the EEG signal 100 ms after the displacement presentation, mainly in occipital and parieto-occipital channels, even in the absence of explicit perception of the displacement.

**Significance Statement:** Stability in vision occurs even while performing saccades. One suggested mechanism for this counterintuitive visual phenomenon is that external displacement is suppressed during the retinal remapping caused by a saccade. Here, we shed light on the mechanisms of trans-saccadic stability by showing that displacement information is not entirely suppressed and specifically present in the early stages of visual processing. Such a signal is relevant and computed for oculomotor adjustment despite being neglected for perception.

## Introduction

Seeing the world requires moving the eyes such that the high-acuity fovea can sample regions of interest that would otherwise remain in the periphery. The benefits of eye movements for vision are undeniable; however, they also pose some problems. Indeed, each eye movement displaces the retinal position of objects, raising the issue of how trans-saccadic perceptual stability is achieved.

Trans-saccadic perceptual stability is studied in the laboratory using gaze-contingent techniques. Participants are asked to report whether a visual target occupies the same spatial location before and after a saccade. They are surprisingly bad at this task, systematically missing displacements up to one-third the amplitude of the saccade, a phenomenon called *saccadic suppression of displacement* (Bridgeman et al., 1975; Deubel et al., 1996). Such observations led scientists to propose that the visual system has a default hypothesis or prior belief about stability (Niemeier et al., 2007). It has been argued that this is efficient given that objects tend to not move during a saccade. Moreover, it prevents the system from mistaking changes in the retinal position of objects due to eye movements for movements in the outside world. Saccade landing positions are generally normally distributed around the visual target, and although some of this variability is centrally planned (Collins et al., 2009), noise arises at lower levels and cannot be predicted (van Beers, 2007). Such a distinction between predicted and unpredicted saccade landing errors is crucial. Indeed, when the saccade does not land on the target but this was predicted (i.e., represented in the efference copy of the saccade), no prediction error occurs. However, in the case of a target displacement, targeting error occurs, but this was not predicted.

Thus, after a saccade, there can be a prediction error between the predicted and actual retinal position of the target, which is not necessarily the same as the target eccentricity. Although it does not lead to the perception of object displacement, it does lead to saccadic adaptation (Collins & Wallman, 2012; McLaughlin, 1967). Saccadic adaptation is the adjustment of saccade amplitude as a function of previous prediction errors. Initially observed in patients recovering from unilateral extraocular muscle paresis, adaptation can be observed in healthy participants by displacing the target during the saccade and observing the evolution of saccade amplitude: backwards displacements lead to amplitude-decreasing adaptation, whereas forwards displacements lead to amplitude-increasing adaptation. Adaptation also occurs on a trial-by-trial basis: an undershoot on one trial tends to be followed by a saccade of greater amplitude on the next. However, when the targeting errors are too large, there is little or no consequence for the next saccade target (Robinson et al., 2003). Moreover, there tends to be a better correlation between targeting error on one trial and amplitude adjustment on the subsequent trial when observers report not seeing the displacement (Collins, 2014).

The targeting errors that do not inform perception do inform the motor system. The present study aimed to identify a brain signature of the prediction errors in the absence of explicit perception of a physical object displacement. Healthy human participants were asked to perform a saccade towards a visual target and to report whether it moved or not during the saccade. We recorded their brain activity via electroencephalography (EEG). We used multivariate pattern analysis to distinguish the brain activity linked to target displacement, a proxy for the prediction error, grounded on the assumption that displacements generate unexpected targeting errors. We performed these analyses separately when observers reported a target displacement and when they reported no displacement. Our primary interest was the no-displacement response trials, since decoding brain activity linked to these trials permits identifying whether and when the brain represents the prediction error when explicit perception is lacking.

## Material and methods

### Data and script availability

All data and analysis scripts will be made freely available online upon publication. The procedures and analysis were predetermined but not pre-registered on a public repository.

### Participants

Nineteen volunteers (aged 25.42 ± 5.05; 10 female) took part in the experiment for an allowance of 15€/hour. All participants had a normal or corrected-to-normal vision and were naive to the hypotheses under investigation. They all gave written informed consent before participating in the experiment. This study was conducted in agreement with the requirements of the Declaration of Helsinki (2004 version) and approved by the local ethics committee (CERES) of Université Paris Cité.

### Apparatus

Stimuli were presented on a CRT Sony GDM-C520 (100 Hz refresh rate) with a resolution of 1024 × 768 pixels and 40 cm width. Stimulus presentation and data collection were performed using Matlab (The MathWorks) with the Psychophysics Toolbox and Eyelink Toolbox extensions (Brainard, 1997; Cornelissen et al., 2002). EEG data were recorded with 64 Ag/AgC1 electrodes mounted on an elastic cap and amplified by an ActiCHamp amplifier (Brain Products). The sampling rate of the EEG recording was set to 500 Hz. Electrodes were arranged according to the international 10-20 systems. Two of the electrodes (TP9 and TP10) were used to record horizontal eye movements (left and right HEOG, respectively), and two other electrodes (FT9 and FT10) were placed on the mastoids (M1 and M2, respectively). The right mastoid (M2) was used as the online reference. Viewing was binocular, and movements of the right eye were monitored with an EyeLink 1000 (SR Research, Mississauga, Ontario, Canada) at a 1000 Hz sampling rate. Head movements were restrained with a chin rest located 51 cm from the screen.

### Main experiment procedure

To start the trial, we instructed the participants to fixate an open black point (0.2 degrees of visual angle, dva, in diameter) displayed at the center of a gray screen (Figure 1A). After 1000 ms of fixation, plus a random jitter between 10 and 40 ms, a target was presented. The target was a vertically oriented Gabor patch (1.7 dva in diameter; 3 cycles) displayed at an eccentricity of 8 dva either to the left or to the right of fixation and at the same vertical level as the fixation dot. When the fixation point disappeared (i.e., after a random interval selected between 360 and 390 ms from target presentation), participants had up to 1000 ms to perform a saccade towards the target. If they performed the eye-movement before or 1000 ms after the go-signal (fixation dot disappearance), the trial was canceled, an error message (‘too early’ or ‘too late’ respectively) appeared on the screen, and a new trial was added to the end of the trial list.

**Figure 1:**
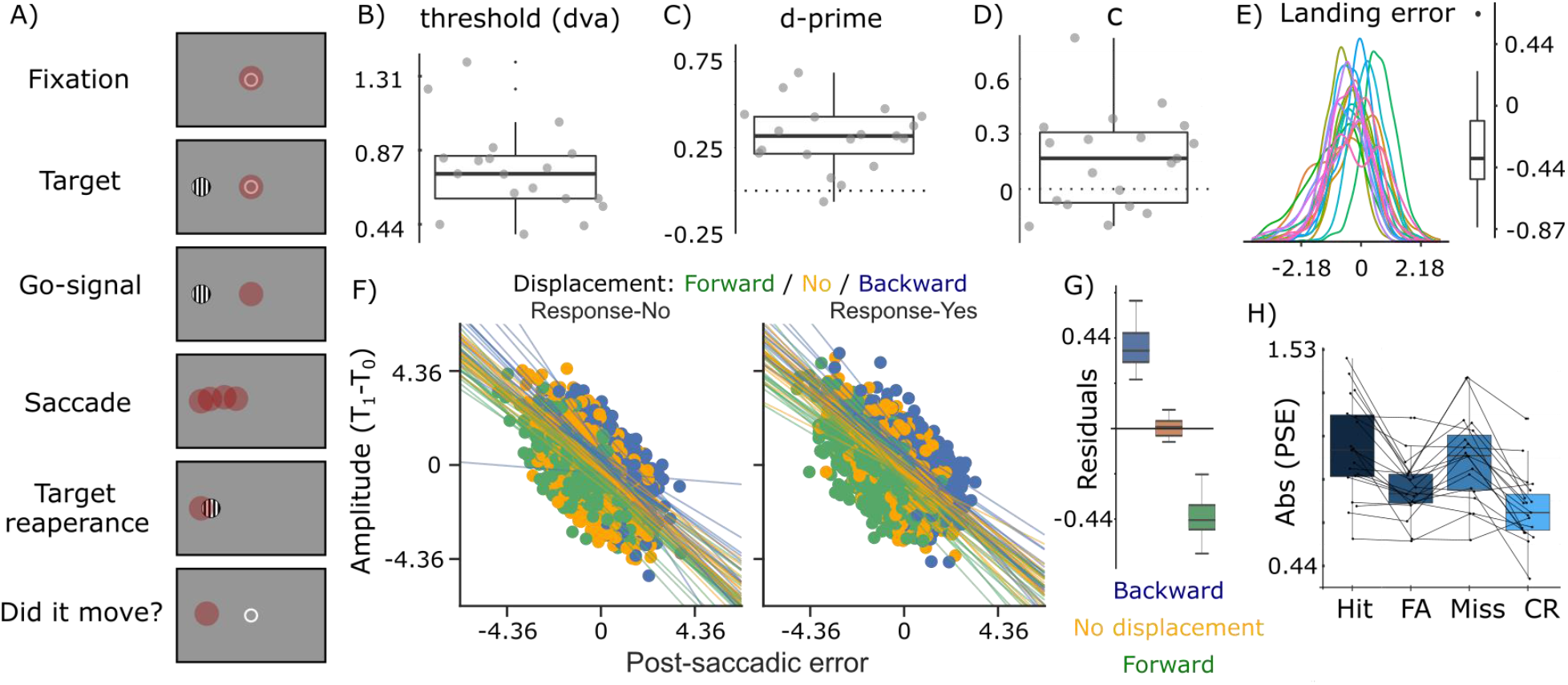
(A) Time course of a single trial. The red dot represents the current eye position. After a go-signal, participants performed a saccade towards a target placed either on the left or on the right of a central fixation point. At saccade onset the target disappeared and reappeared 50 ms after at the same place (no displacement, 50%) or at a distance *d* away from the initial position (displacement condition, 25% backward, 25% forward). Participants reported whether the target moved or not. (B) Average distance thresholds *d* in degrees of visual angle measured in the staircase task. (C) Average sensitivity d’ to target displacements. (D) Decision criterion. From panels B to D, all averages were calculated across participants, and each point represents an individual observer. (E) Individual distributions of landing errors and the boxplot of median error across participants. Negative values represent undershoots and positive values overshoots. (F) Relationship between post-saccadic landing error and adjustment in subsequent saccadic amplitude. Lines represent their linear relationship for each participant, displacement condition (Backward / No displacement / Forward) and response type (No / Yes). (G) Median amplitude residuals of the linear relationship of saccadic adjustment and post-saccadic landing errors separated by displacement condition (backward, no-displacement, forward) across participants. Post-saccadic errors were not enough for explaining the saccadic adjustment since the actual adjustment is below/above the expected in the forward/backward conditions respectively. This intuition can be extracted from panel I as well, where at no adjustment (y=0), participants systematically had more positive errors (overshoots) in backward trials than in no-displacement trials, and more negative errors (undershoots) in forward trials. (H) Distribution of the median absolute post-saccadic error for hits, false alarms, misses and correct rejections across participants.

A saccade was considered to have been initiated when gaze direction moved 2° away from the center of fixation. Once a saccade was detected, the target disappeared. It reappeared 50 ms later, either at the same location (on 50% of the trials), or at some distance *d* to the left or right of the initial target location. The 50 ms blank period between the action and the post-saccadic target presentation was introduced to decrease the overlapping of saccade and target-displacement EEG responses. The target was displayed for an additional 600 ms and then disappeared. After it disappeared, participants had to respond whether they believed the target had moved or not from its initial position. They used the arrow key ‘up’ to denote *“yes, the target moved*”, and the ‘down’ key for *“no, the target did not move”*. Responses were reported with the index finger of the right hand. After their response, participants had 1 second to blink before a new trial began. A break was proposed every 32 trials during which the experimenter was able to perform an eye-tracker calibration if necessary.

The distance *d* was set individually using a staircase procedure (described below) that each participant completed before the main experiment at the beginning of the session. More specifically, the distance *d* was adjusted to lead to an average performance accuracy ranging between 50 and 60%.

A series of operational errors could lead to the trial being canceled and added to the end of the trial list. This was the case if participants’ saccade failed to land within 4 dva from the target’s initial position (a radius of half of target horizontal eccentricity), if participants responded before the target disappeared, or if they blinked at any moment between the beginning of the trial and the disappearance of the post-saccadic target.

The experiment ended when participants had performed a total of 640 valid trials (i.e., without any of the operational errors described above) resulting from 160 repetitions of the combinations of 2 displacements (present, absent) and 2 saccade directions (left, right). The task was slightly shorter for the first two participants who completed 512 trials (2 displacements (present, absent) * 2 saccade directions (left, right) * 128 trials). Only valid trials were analyzed.

### Staircase procedure

Before the main experiment, participants ran a staircase procedure. The task was identical to the one described above, except that during the staircase, no EEG data were recorded, and participants were required to report whether the target moved to the left or the right of its initial location using the ‘left’ or ‘right’ arrow key, respectively. We did not include trials with no displacement, and the direction (left or right) was selected randomly. Three interleaved staircases controlled the amplitude of the left/right target displacement. Each staircase started with a different displacement amplitude, i.e., 0.5, 1, and 2 dva. If the observer’s response was correct (incorrect), the amplitude of the displacement (left or right) decreased (increased) in the subsequent trial. The magnitude of the displacement was controlled by an accelerated stochastic approximation algorithm (Kesten, 1958) set to converge at the displacement amplitude leading to an average performance accuracy ranging between 50% and 60% correct responses. Each staircase stopped once the convergence level was reached. The convergence values obtained with the three staircases were averaged and used in the main experiment trials when a target movement was displayed. The staircase procedure lasted approximately 20 min.

### Behavioral analysis

Trials were divided based on the combination of participants’ judgment (yes, no) and target displacement (displacement, no displacement), leading to four experimental conditions corresponding to the four possible outcomes of Signal Detection Theory: Hits (observers reported that the target moved and the target had indeed moved), False Alarms (observers reported that the target moved but the target had not moved), Correct Rejections (observers reported that the target did not move and the target indeed had not moved) and Misses (observers reported that the target did not move but the target had moved). Participants’ performance was quantified by calculating detection sensitivity *d’* and decision criterion *c*.

Saccadic landing errors were calculated as the difference between the eye position when the target reappeared (i.e., 50 ms after the onset of the saccade) and the target’s initial position. In addition, the sign of the error was flipped for the left saccade trials, such that positive errors expressed overshooting saccades and negative errors undershooting saccades.

The post-saccadic error was calculated as the difference between the eye position when the target reappeared and the final target position. To assess saccadic adaptation, we computed the change in saccadic amplitude between successive trials. More specifically, the saccadic amplitude at trial *t1* was subtracted from the amplitude observed at trial *t0*. Then we fitted a generalized linear model (GLM), implemented in statsmodels library (v.0.10.1) in Python for each participant to predict the saccadic adjustment. The function took into account the following factors: the post-saccadic error, displacement (−1 for backward, 0 for no-displacement, 1 for forward), response (0 for no-displacement, 1 for displacement), and interactions. Saccade amplitude and post-saccadic error were expressed in pixels. The statistical significance of the factors was evaluated at the group level via a one-sample t-test (μ = 0, α = 0.05).

### EEG analyses

#### Preprocessing

Preprocessing of continuous EEG activity was conducted in Python (v.3.6.9) by using the MNE (v.0.18.2) toolbox (Gramfort et al., 2014). Data were re-referenced to the average of left and right mastoids and then filtered with a 4th order Butterworth non-causal filter between 0.5 to 30 Hz. Data were segmented from -200 ms to 600 ms relative to saccade onset. EEG segments were then corrected with a 200 ms baseline time-locked to the first target presentation in the trial (i.e., from -200 ms to 0 ms relative to target onset).

#### Multivariate pattern analysis

EEG data were analyzed using MultiVariate Pattern Analyses (MVPA) performed in Python using the Scikit-learn library (v.0.22.1). Different models were applied depending on the analyzed signal (EEG or Eye-tracking data).

We were particularly interested in evaluating whether we could classify displacement and no displacement target trials from brain activity. One model aimed at classifying displacement and no-displacement trials when participants reported a target displacement (i.e., hit versus false alarm trials), and another model aimed at dissociating the same trials but when participants did not report experiencing a displacement (i.e., correct rejection versus miss trials). Importantly, to ensure that classification accuracy was not driven by eye-movements occurring after saccade landing, MVPA were performed on all trials, excluding those that contained micro-saccades occurring within a time window going from 50 to 200 ms after the reappearance of the target (see the section “Eye-position control analyses” below for more details).

Trial-by-trial EEG classification was performed with an L2-regularized logistic classifier with a *liblinear* solver. Features consisted of the EEG activity recorded from all electrodes, except the references and HEOG electrodes. Feature scaling was performed as a preprocessing step, namely, data were normalized (z-score transformation) for each time point across trials, using the parameters (mean and standard deviation) of the training set. Our classification procedure implemented a stratified 5-fold cross-validation approach for each time point of the segmented data. Scores were the area under the receiver operating characteristic curve (AUC), which we calculated for each fold and reported their average.

For each time point, we computed the weight vectors from each linear model to assess the electrodes that contributed the most to classification accuracy. Later, they were transformed into activation patterns as implemented in MNE to provide a neurophysiological interpretation to the coefficients, implying that electrodes with nonzero values carry class-specific information (Haufe et al., 2014).

Finally, to assess significant classification from chance (AUC = 0.5) at the participant level, we used cluster-based permutation tests based on paired t-scores (Maris & Oostenveld, 2007), with 10000 random permutations, alpha value as 0.05, implemented in MNE by the function *permutation_cluster_1samp_test*. In addition, for the evaluation of activation maps, we performed permutation t-tests on electrodes’ data as implemented in MNE by the function *permutation_t_test*, with 10000 random permutations and alpha as 0.0125 (alpha of 0.05 divided by four time-window tests).

### Control analyses

#### Eye position

Even in the absence of conscious perception of target displacement, volunteers could move their gaze towards the displaced target, thus potentially contaminating the EEG signal. Therefore, we removed (offline) the trials containing (micro)saccades within a time window ranging from 50 to 200 ms after the main saccade detection (proportion of excluded trials = 0.044 ± 0.062). We selected this time window since it exhibited the largest EEG decoding accuracy (see Results section for more details). Microsaccades were identified using the Microsaccade Toolbox 0.9 implemented in R, considering the eye tracking sampling rate (1000 Hz) and 6 ms as the minimum duration criteria and velocity threshold scaling factor as 5. The MVPA on EEG data was performed on this set of trials.

Furthermore, to ensure that EEG decoding accuracy was not merely driven by eye movements, we conducted classification analyses to dissociate displacement from no-displacement trials using gaze position data. Gaze position data corresponded to the online horizontal and vertical eye position, which was recorded throughout the experiment. Specifically, as for the EEG data, we performed the classification of the displacement separately for the responses *“Yes, the target moved”* (i.e., hit versus false alarms trials), and *“No, the target did not move”* (i.e., correct rejection versus miss trials). We used a non-linear classifier (C-Support Vector Classification with RBF kernel and gamma as auto) because better classification boundaries are generated as the target could move to two different directions (see Figure 3B). Features consisted of the gaze position in the vertical axis and the gaze’s absolute distance from the center of the screen in the horizontal axis. This transformation in the x-axis enabled us to mirror the left hemifield and aggregate gaze position of the left and right trials. Data were normalized using z-score transformation and classification was performed with a stratified 5-fold cross-validation approach for each time point. Classification accuracy was summarized by the area under the receiver operating characteristic curve (AUC).

#### Target eccentricity

To evaluate a possible role of saccadic amplitudes and post-saccadic error (i.e., the distance between the new target position and gaze location after the saccade) on the EEG decoding, the same EEG-MVPA protocol described above was implemented with the exception that classifiers were trained and tested on the activity of parieto-occipital electrodes (O1, Oz, O2, PO7, PO3, POz, PO4, PO8), and within a time window going from 100 to 200 ms after the saccade. Instead of averaging the activity in time for each electrode, we used the signal at each time point as an attribute, summing up to 408 independent variables (8 electrodes x 51 time points). We focused on parieto-occipital channels since these electrodes were more strongly responsible for EEG decoding accuracy (see *Results* section for more details). Then, for each participant and each response model, we fitted the predicted EEG classifier labels (1 for displacement and 0 for no-displacement) with a Binomial GLM which included saccadic amplitude, target eccentricity (a.k.a. absolute post-saccadic error) and their interaction as predictors. The statistical significance of *beta* values was assessed, at the group level, with a one-sample t-test (μ = 0, α = 0.05) for each response-model.

#### Behavioral relevance of EEG decoding

In a second step, we aimed to verify whether the prediction error signal present in parieto-occipital electrodes was a variable explaining the saccadic amplitude adjustment. Thus, we fitted mixed models and calculated Akaike’s Information Criteria (AIC) and Bayesian Information Criteria (BIC) to compare their goodness of fit. The mixed-models and goodness of fit calculations were performed with the statsmodels library (v.0.10.1).

The first mixed-model (null-model) considered the post-saccadic error and response in trial *t0* as fixed factors (with interactions) and participants as the random factor to model the amplitude adjustment in trial *t1*. A second mixed model (full model) considered the post-saccadic error, the actual target *displacement* (backward, forward and no displacement), and response as fixed factors (with interactions). The factor “*participants”* was included as a random intercept. The third and fourth models were identical to the second, except that rather than using the actual target displacement to predict saccadic adaptation, we used the target displacement labels predicted by the EEG decoders described below.

The same EEG-MVPA method reported in the section “Multivariate pattern analysis” was performed with two exceptions. First, classifiers were trained and tested on the activity of parieto-occipital electrodes (O1, Oz, O2, PO7, PO3, POz, PO4, PO8) observed within a time window going from 100 to 200 ms after the saccade, corresponding to 408 independent variables (8 electrodes x 51 time points). Second, the one-versus-rest approach was selected to decode backward displacement, forward displacement, and no displacement trials. For the third mixed model (*true decoding model*), the displacement labels used to predict saccadic adaptation were the *good* guesses of the *true* classifier, i.e., the classifier exposed to the proper labels during training. Instead, the fourth mixed model (*shuffled decoding model*) used the displacement labels predicted by a classifier trained on shuffled labels. In other words, for the shuffled classifier, we removed the actual correspondence between brain activity and actual target displacement. The model predicting saccadic adaptation from the output of the shuffled decoder was compared with the true decoding model by calculating the AIC and BIC estimators, in order to evaluate whether the good guesses of the true decoder could predict saccadic adaptation. The AIC and BIC criteria favor models with the closest probability distribution to the fitted data while controlling for overfitting issues. In fact, simply adding a random parameter to a model may increase its likelihood. However, AIC and BIC attribute penalties for additional parameters (the penalty term is larger in BIC than AIC). Therefore, we expected the shuffled decoder to be the worst model.

## Results

### Behavioral results

The individual displacement distances ranged from 0.38 to 1.39 dva (Q_1/4_ = 0.59, Q_2/4_ = 0.73, Q_3/4_ = 0.85, Figure 1B), and as expected by the experimental design, participants exhibited low sensitivity *d’* to target displacement (Figure 1C). They also showed positive decision criterion values indicating a tendency towards a conservative response bias (Q_1/4_ = -0.08, Q_2/4_ = 0.17, Q_3/4_ = 0.32, Wilcoxon signed-rank test, Z = 2.374, p = 0.018, Figure 1D). Additionally, the hit rates for forward (outward) and backward (inward) displacements were not significantly different (Hit rate (mean ± std) for backward = 0.512 ± 0.237 and forward = 0.456 ± 0.185; t(18) = -0.048, p = 0.962). Overall, participants performed the task near their perceptual threshold (Proportion of correct responses: Q_1/4_ = 0.54, Q_2/4_ = 0.56, Q_3/4_ = 0.58).

Saccades ended 15 ms (median time) after the target disappearance, so in most of the trials there was no target when the participants’ eyes landed. The analysis of saccade landing errors showed that saccades undershot with respect to the initial position of the target (Figure 1E). Notwithstanding, participants adapted their saccadic performance on a trial-by-trial basis. Post-saccadic errors (pse) resulting from saccade undershoots led to a saccade with larger amplitude in the following trial, whereas saccade overshoots resulted in subsequently smaller saccadic amplitudes (β mean ± std = -0.865 ± 0.094, t(18) = -40.189, p < 0.001, Figure 1F). This replicates previous work, except that we did not find a modulation in saccadic adaptation by participants’ response (response: β = -0.168 ± 3.181, t(18) = -0.231, p = 0.82; pse-response interaction: β = 0.017 ± 0.099, t(18) = 0.734, p = 0.472; response-displacement interaction: β = 0.002 ± 3.386, t(18) = 0.003, p = 0.998; pse-response-displacement interaction: β = -0.014 ± 0.136, t(18) = -0.448, p = 0.66). Saccadic adaptation also depended on the target displacement (β = -14.577 ± 6.767, t(18) = -9.389, p < 0.001, Figure 1G). Notably, when participants were presented with a backward (forward) displacement, they tended to execute larger (smaller) saccades in the subsequent trial. However, the actual adjustment was often smaller than the adjustment one would expect from the forward/backward conditions (e.g., if in a given trial the target was displaced by 1 dva backward or forward, in the subsequent trial the adjustment was smaller than 1 dva).

Finally, we observed no interaction between post-saccadic errors and displacement (pse-displacement: β = 0.031 ± 0.087, t(18) = 1.557, p = 0.137), i.e., the strength of saccadic adaptation was comparable across the different displacement conditions (backward: pse β = -0.878 ± 0.123, no-displacement: pse β =-0.832 ± 0.076, forward: pse β = -0.846 ± 0.127; paired t-test backward-forward: t(18) = -0.994, p = 0.334; paired t-test backward-no: t(18) = -1.543, p = 0.14; paired t-test forward-no: t(18) = -0.534, p = 0.6).

### Unseen post-saccadic displacement is decodable at early stages of visual processing

Despite low sensitivity to target displacements, participants nevertheless adapted their saccades based on the post-saccadic displacement and errors. Therefore, we investigated whether displacement information was present in the EEG signal. We were particularly interested in whether we could decode displacement information (i.e., the target moved vs the target did not move) in the cases in which participants replied *“no, the target did not move”* since these unseen displacements still led to trial-by-trial adaptation. Although classification score was not very high (AUC < 0.6), we could decode target displacement above chance (Figure 2) both when participants reported that the target moved (cluster-based permutation tests: from 166 to 212 ms, p = 0.026; from 298 to 348 ms, p = 0.02; from 362 to 416 ms, p = 0.021; 498 to 552 ms, p = 0.028) and when they reported that it did not move (cluster-based permutation tests, no-response: from 128 to 338 ms, p < 0.001). Classification scores along time were comparable for the response models (cluster-based permutation test, p = 0.938 and p = 0.482).

**Figure 2:**
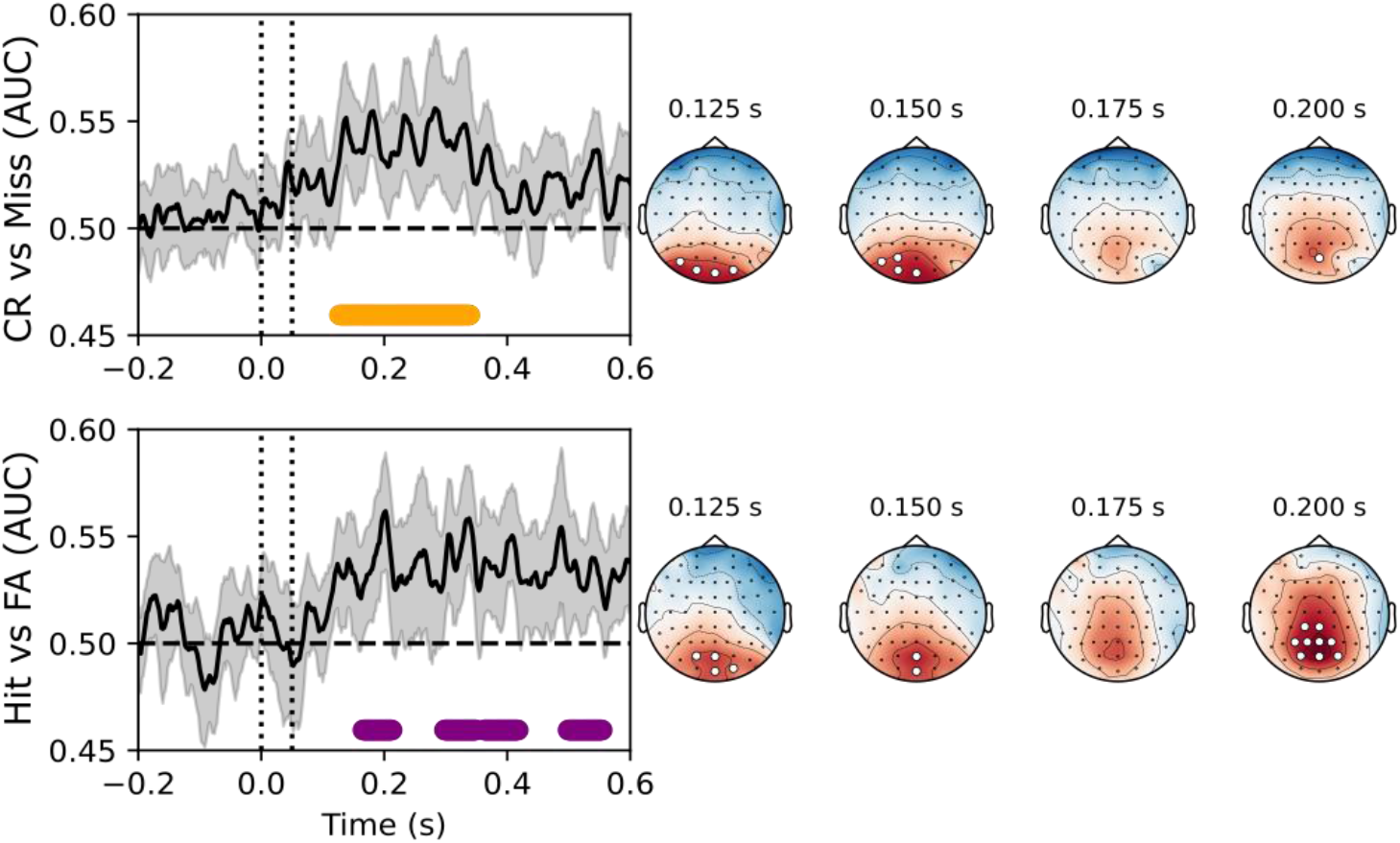
Decoding displacements from the EEG signal. Dotted vertical lines represent saccade detection (at 0 s) and target reappearance (at 0.05 s). Bold dots at the bottom of the panels represent time points at which classification performance differs from chance (AUC = 0.5). Right: topographic activation maps at four time windows, with the significant channels highlighted. The colormap ranges from -10 (cool colors) to 10 (warm colors) arbitrary units.

**Figure 3:**
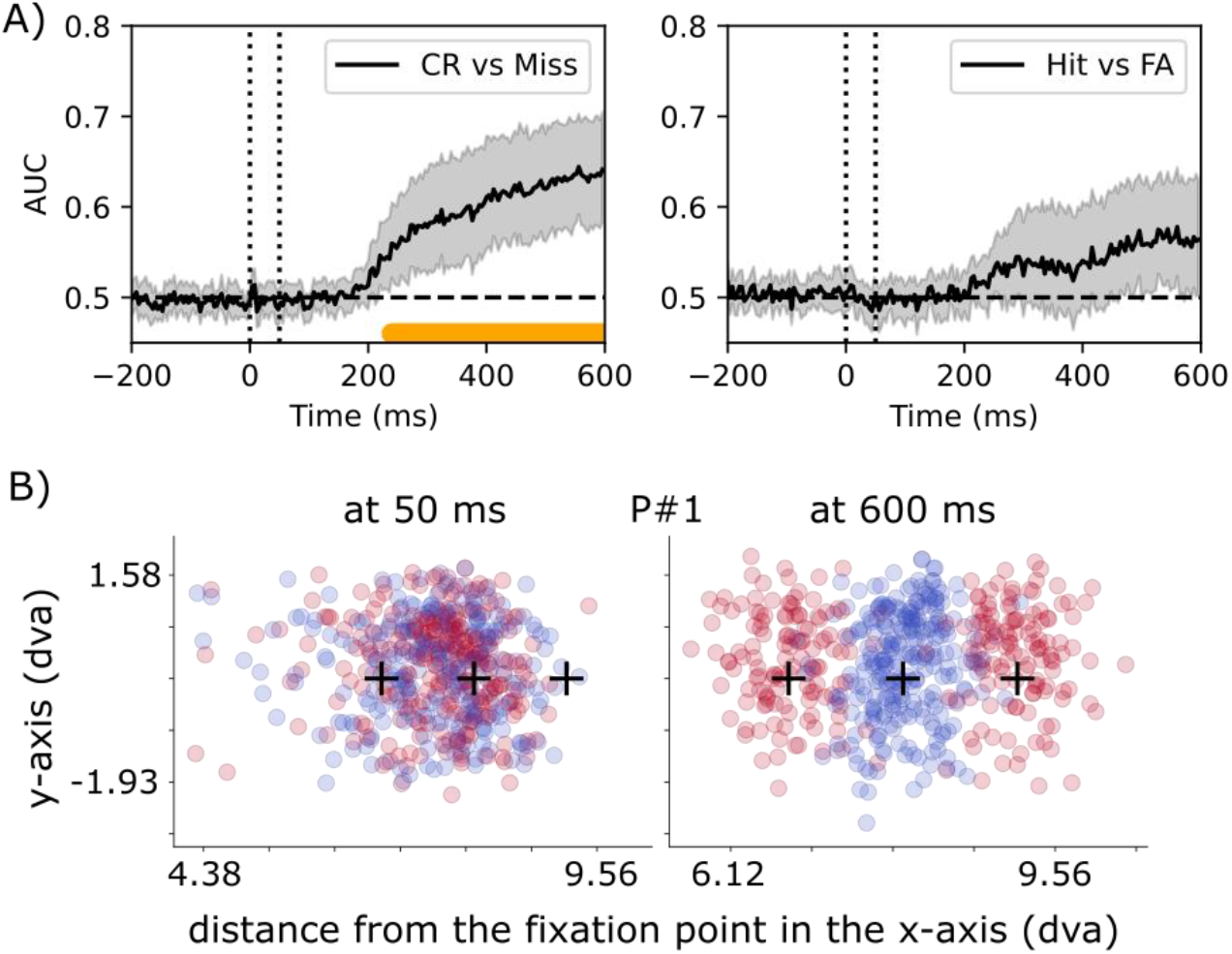
(A) Decoding displacements from the eye position data. Dashed lines and shaded areas as in Figure 2. (B) Eye position data of participant #1 at 50 ms (target reappearance) and at 600 ms (right before the response go-signal). The y-axis is the vertical position on the screen, while the x-axis is the distance from the center of the screen, the initial fixation. The plus sign markers represent the three possible positions of the target across trials (backward displacement, no-displacement, forward displacement). The gaze of participant #1 in all trials are represented by the dots. The dot color represents whether there was a displacement (in red) or not (in blue) in each trial. As in A, it shows that for participant #1, there is a clear boundary for classifying the displacement from the eye position at the end of the trial (600 ms), but not at the beginning when the target just reappeared (50 ms), meaning that participant #1 generally performed eye movements towards the target at some point of the trial.

To understand which electrodes were contributing the most to the significant classification from 100 to 200 ms, we analyzed the activation patterns generated by the linear classifiers at four time windows within this period (115-135 ms; 140-160 ms; 165-185 ms, 190-210 ms). The electrodes that contributed the most to displacement classification were occipital and parieto-occipital electrodes (Permutation tests; response-yes: 115-135 ms = Oz, O2, PO3, POz; 140-160 ms = Oz, POz; 190-210 ms = CP1, Pz, P3, P1, PO3, POz, PO4, P2, CPz; response-no: 115-135 ms = O1, Oz, O2, PO7; 140-160 ms = O1, Oz, PO7, PO3; 190-210 ms = POz). This suggests that information about post-saccadic displacement is represented in cortical visual areas during the early visual processing stages, even when this information will not subsequently lead to a perception of displacement.

Post-saccadic displacement classification during early processing stages (100-200 ms) cannot be explained by eye movements since we excluded trials with microsaccades within this window. Additionally, when we performed MVPA but using the gaze position data, scores were only different from chance from 200 ms onwards (Figure 3A; cluster-based permutation tests: response-no: p < 0.001, response-yes: p = 0.363), suggesting that saccade landing positions were similar across conditions, but after 150 ms the target reappearance in displaced trials participants start to correct their gaze towards the target (Figure 3B). All considered, this confirms that the early decoding of displacement is not due to eye movements.

Another alternative explanation is that the significant classification was not related to an error-related signal but due to differences in post-saccadic errors (i.e., the distance between the new target position and gaze location after the saccade). Indeed, by design, *displacement* trials generated bigger post-saccadic errors than *no-displacement* trials (see Figure 1H). One way of dealing with this confound would be to consider comparable parts of the distribution of errors between conditions, but of course this would create a new confound because those trials would no longer be comparable with regards to saccadic amplitude. For this reason, a series of binomial GLM investigated whether error size and saccadic amplitude could predict the displacement labels resulting from EEG classifiers (for details see the Control analyses session). The rationale was to verify whether the size of the post-saccadic error and/or the saccadic amplitude were the variables explaining the label assignment. If this was the case, then we would expect for example that small post-saccadic errors would be consistently associated with no-displacement labels and bigger errors with displacement labels.

Additionally, linear mixed models assessed whether the target displacement labels predicted by the EEG classifiers could predict saccadic adaptation over and above post-saccadic errors. Taken together these analyses would highlight whether such brain activity contained specific information about trans-saccadic prediction error that was not merely reflected by the size of post-saccadic errors. As indicated above, for these analyses, classifiers were trained and tested on the activity of parieto-occipital electrodes (i.e., the electrodes most responsible for decoding accuracy) during the time window going from 100 to 200 ms after the saccade (i.e., a time window in which EEG signals were not contaminated by eye-movements).

Displacement classification remained significant using the signal from parieto-occipital channels during early time points (Response-yes: AUC = 0.571 ± 0.064, t(18) = 4.802, p < 0.001; Response-no: AUC = 0.574 ± 0.093, t(18) = 3.505, p = 0.003). Critically, the size of post-saccadic errors and saccadic amplitude did not predict the displacement labels provided by the EEG classifier. This was the case both in trials in which participants responded *target moved* (error size: β = -0.024 ± 0.08, t(18) = -1.31, p = 0.207; amplitude: β = 0 ± 0.018, t(18) = -0.083, p = 0.935; interaction: β = 2.03e-04 ± 4.45e-04, t(18) = 1.957, p = 0.066) and *target did not move* (error size: β = 0.024 ± 0.14, t(18) = 0.761, p = 0.457; amplitude: β = 0.002 ± 0.021, t(18) = 0.456, p = 0.654; interaction: β = -6.14e-05 ± 7.75e-04, t(18) = -0.346, p = 0.734). Similar results are seen when performing this analysis but on the subset of trials that were correctly classified, with the exception that in this case the interaction of saccadic amplitude and error size was significant in Response-yes trials (β = -0.001 ± 0.003, t(18) = 2.137, p = 0.047). We can thus exclude that post-saccadic target eccentricity and saccadic amplitude were the information source for displacement EEG classification. We do not claim that occipital and parietal activity does not reflect visual post-saccadic target representation, but our results suggest that it also reflects a representation of prediction error.

The following analyses investigated whether the displacement labels resulting from the EEG classification predicted saccadic adaptation, consequently, representing prediction errors. We fitted four mixed models for the amplitude adjustment and compared their goodness-of-fit. The null model included post-saccadic errors and response type as predictors. The other three models nested the null-behavioral model plus a displacement variable. In the full-behavioral model the displacement variable consisted of the actual target displacement observed in the trials. In the true decoding model, this variable contained the displacement labels predicted by the classifiers trained with the actual target displacement. Finally, in the shuffled decoding model, the displacement variable consisted of the displacement labels predicted by the shuffled decoders (i.e., these classifiers were trained on a set of trials in which we shuffled the actual target displacement labels). Table 1 summarizes models’ AIC and BIC values and shows again by another method that the full behavioral model had the best fit. The second best model was the true decoding model, with lower AIC and BIC values than the null and shuffled decoding model. Additionally, we ran likelihood ratio tests comparing each full model against the null behavioral model. Once more the analyses showed that displacement variables significantly improved the fit (full behavioral model: χ^2^(4) = 2206.03, p < 0.001; true decoding model: χ^2^(4) = 77.66, p < 0.001; shuffled decoding: χ^2^(4) = 19.452, p = 0.001). In sum, these analyses showed that the signal observed from 100 to 200 ms in parieto-occipital sites contains information that is not merely reflected by post-saccadic errors, and that is relevant for predicting the future saccadic behavior.

**Table 1:**
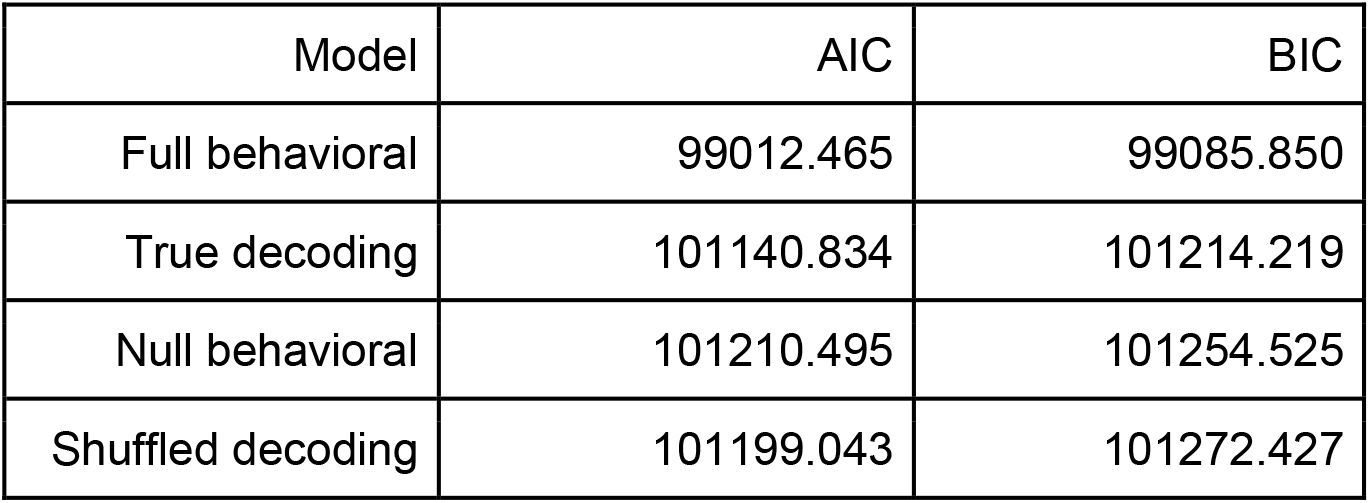
Goodness-of-fit measured by Akaike’s Information Criteria (AIC) and Bayesian Information Criteria (BIC) for the estimated mixed-models. Smaller values correspond to better fits.

| Model | AIC | BIC |
| --- | --- | --- |
| Full behavioral | 99012.465 | 99085.850 |
| True decoding | 101140.834 | 101214.219 |
| Null behavioral | 101210.495 | 101254.525 |
| Shuffled decoding | 101199.043 | 101272.427 |

## Discussion

This study investigated the brain signatures of trans-saccadic prediction errors in the absence of the conscious perception of target displacement. Participants performed a saccade toward a visual target which disappeared when the saccade was initiated and reappeared 50 ms later either in the same or in a different location. We decoded the presence versus absence of displacement both when participants were aware of a displacement and when they were not. We argue that the signal driving EEG decoding accuracy is the prediction error between the expected and observed saccade landing position, and that this signal is present at early stages of processing, even when it drives motor learning but does not inform perception.

Saccadic suppression of displacement (SSD) refers to the absence of awareness of trans-saccadic displacements (Bridgeman et al., 1975). Such blindness was taken as evidence that the visual system discards or ignores these displacements. However, two findings show that this blindness is not due to low sensitivity. First, performance is near perfect when the post-saccadic target reappears at its displaced location several hundred milliseconds after the saccade has landed (Deubel et al., 1996, 1998). Second, saccade amplitude changes trial by trial as a function of these “unseen” displacements, suggesting that while they are ignored by explicit perception, they are not ignored by the motor system, which continually adjusts to make up for what it considers to be its own errors (Collins, 2014). We replicated this trial-by-trial saccadic adaptation in the current data set. We further showed that the motor system has access to information about target displacement at early stages during visual processing. Indeed, EEG classifiers could dissociate displacement and no-displacement trials from about 100 ms after saccade onset. Moreover, occipital and parieto-occipital channels were the electrodes mostly responsible for classification accuracy.

These results are compatible with previous studies reporting an involvement of parietal areas in error signals related to saccades (Collins & Jacquet, 2018). Such error signals are thought to represent a discrepancy between the predicted landing position and the observed landing position. We believe that the post-saccadic displacement and error that we measured here reflect such an error signal. Neuropsychological studies reported deficits in saccade tasks compatible with the involvement of posterior parietal cortex (PPC) in saccade-related error signals (Duhamel et al., 1992; Gaymard et al., 1994; Heide & Kömpf, 1998; Pisella et al., 2011), but the often large extent of the lesions and the relatively small sample sizes make it difficult to pinpoint a more specific locus. Imaging studies also point towards the involvement of the PPC in saccade-related error signals: PPC has an eye-centered organization that is updated when the eyes move (Bellebaum et al., 2005; Medendorp et al., 2003) according to saccadic goals and not to the coordinates of the visual stimulus (Medendorp et al., 2005).

Furthermore, some PPC neurons show predictive remapping of their receptive fields: just prior to a saccade, receptive fields become responsive to stimuli in the future retinotopic location (Duhamel et al., 1992; Hall & Colby, 2011). This means that the PPC receives information about saccade error and updates retinotopic visual maps based on this signal. The expected oculomotor error is transmitted via efference copy and is available approximately 30 ms before the onset of a saccade. It has been shown that this prediction is usually correct (Collins et al., 2009) and that saccadic adaptation depends mostly on its prediction error (Collins & Wallman, 2012).

Our interpretation of the EEG decoding relies on the assumptions that saccadic adaptation is driven by prediction error and that internal prediction of saccadic landing positions is usually correct such that prediction errors only occur when the target is displaced. Our interpretation would need to be revised if, for example, saccadic adaptation resulted from postdictive updates of space rather than prediction errors (Masselink & Lappe, 2021). The alternative explanation that adaptation would be due to post-saccadic target information (Bellebaum et al., 2005) cannot explain our results because we explicitly showed that decoding labels were not predicted by post-saccadic landing positions. Furthermore, EEG decoding labels provided relevant information that improved the prediction of saccadic adaptation. In sum, the prediction error account of the SSD phenomenon seems to best fit our results.

All things considered, our results are based on current assumptions about visuomotor learning and coherent with these studies in that we were able to decode unseen displacements successfully. However, our results also go further than previous work showing that saccade-related error signals are also present in the early stages of visual processing.

## Acknowledgments

This research was funded by the Agence Nationale de la Recherche, research funding ANR JCJC, project number ANR-18-CE10-0001, awarded to Andrea Desantis. We are grateful to Dr Qing Yang for his help in the data collection of the current experiment.

## References

Bellebaum, C., Hoffmann, K.-P., & Daum, I. (2005). Post-saccadic updating of visual space in the posterior parietal cortex in humans. Behavioural Brain Research, 163(2), 194–203.

Brainard, D. H. (1997). The Psychophysics Toolbox. In Spatial Vision (Vol. 10, Issue 4, pp. 433– 436). https://doi.org/10.1163/156856897×00357

Bridgeman, B., Hendry, D., & Stark, L. (1975). Failure to detect displacement of the visual world during saccadic eye movements. Vision Research, 15(6), 719–722.

Collins, T. (2014). Trade-off between spatiotopy and saccadic plasticity. Journal of Vision, 14(12). https://doi.org/10.1167/14.12.28

Collins, T., & Jacquet, P. O. (2018). TMS over posterior parietal cortex disrupts trans-saccadic visual stability. Brain Stimulation, 11(2), 390–399.

Collins, T., Rolfs, M., Deubel, H., & Cavanagh, P. (2009). Post-saccadic location judgments reveal remapping of saccade targets to non-foveal locations. Journal of Vision, 9(5), 29.1–9.

Collins, T., & Wallman, J. (2012). The relative importance of retinal error and prediction in saccadic adaptation. Journal of Neurophysiology, 107(12), 3342–3348.

Cornelissen, F. W., Peters, E. M., & Palmer, J. (2002). The Eyelink Toolbox: Eye tracking with MATLAB and the Psychophysics Toolbox. In Behavior Research Methods, Instruments, & Computers (Vol. 34, Issue 4, pp. 613–617). https://doi.org/10.3758/bf03195489

Deubel, H., Bridgeman, B., & Schneider, W. X. (1998). Immediate post-saccadic information mediates space constancy. Vision Research, 38(20), 3147–3159.

Deubel, H., Schneider, W. X., & Bridgeman, B. (1996). Postsaccadic target blanking prevents saccadic suppression of image displacement. Vision Research, 36(7), 985–996.

Duhamel, J. R., Colby, C. L., & Goldberg, M. E. (1992). The updating of the representation of visual space in parietal cortex by intended eye movements. Science, 255(5040), 90–92.

Gaymard, B., Rivaud, S., & Pierrot-Deseilligny, C. (1994). Impairment of extraretinal eye position signals after central thalamic lesions in humans. Experimental Brain Research. Experimentelle Hirnforschung. Experimentation Cerebrale, 102(1), 1–9.

Gramfort, A., Luessi, M., Larson, E., Engemann, D. A., Strohmeier, D., Brodbeck, C., Parkkonen, L., & Hämäläinen, M. S. (2014). MNE software for processing MEG and EEG data. NeuroImage, 86, 446–460.

Hall, N. J., & Colby, C. L. (2011). Remapping for visual stability. In Philosophical Transactions of the Royal Society B: Biological Sciences (Vol. 366, Issue 1564, pp. 528–539). https://doi.org/10.1098/rstb.2010.0248

Haufe, S., Meinecke, F., Görgen, K., Dähne, S., Haynes, J.-D., Blankertz, B., & Bießmann, F. (2014). On the interpretation of weight vectors of linear models in multivariate neuroimaging. NeuroImage, 87, 96–110.

Heide, W., & Kömpf, D. (1998). Combined deficits of saccades and visuo-spatial orientation after cortical lesions. Experimental Brain Research. Experimentelle Hirnforschung. Experimentation Cerebrale, 123(1-2), 164–171.

Kesten, H. (1958). Accelerated Stochastic Approximation. In The Annals of Mathematical Statistics (Vol. 29, Issue 1, pp. 41–59). https://doi.org/10.1214/aoms/1177706705

Maris, E., & Oostenveld, R. (2007). Nonparametric statistical testing of EEG- and MEG-data. Journal of Neuroscience Methods, 164(1), 177–190.

Masselink, J., & Lappe, M. (2021). Visuomotor learning from postdictive motor error. eLife, 10. https://doi.org/10.7554/eLife.64278

McLaughlin, S. C. (1967). Parametric adjustment in saccadic eye movements. In Perception & Psychophysics (Vol. 2, Issue 8, pp. 359–362). https://doi.org/10.3758/bf03210071

Medendorp, W. P., Goltz, H. C., Crawford, J. D., & Vilis, T. (2005). Integration of target and effector information in human posterior parietal cortex for the planning of action. Journal of Neurophysiology, 93(2), 954–962.

Medendorp, W. P., Goltz, H. C., Vilis, T., & Crawford, J. D. (2003). Gaze-centered updating of visual space in human parietal cortex. The Journal of Neuroscience: The Official Journal of the Society for Neuroscience, 23(15), 6209–6214.

Niemeier, M., Douglas Crawford, J., & Tweed, D. B. (2007). Optimal inference explains dimension-specific contractions of spatial perception. In Experimental Brain Research (Vol. 179, Issue 2, pp. 313–323). https://doi.org/10.1007/s00221-006-0788-9

Pisella, L., Alahyane, N., Blangero, A., Thery, F., Blanc, S., & Pelisson, D. (2011). Right-hemispheric dominance for visual remapping in humans. Philosophical Transactions of the Royal Society of London. Series B, Biological Sciences, 366(1564), 572–585.

Robinson, F. R., Noto, C. T., & Bevans, S. E. (2003). Effect of visual error size on saccade adaptation in monkey. Journal of Neurophysiology, 90(2), 1235–1244.

van Beers, R. J. (2007). The sources of variability in saccadic eye movements. The Journal of Neuroscience: The Official Journal of the Society for Neuroscience, 27(33), 8757–8770.

